# Evolution of antivirus defense in prokaryotes depending on the environmental virus prevalence and virome dynamics

**DOI:** 10.1101/2025.05.27.656525

**Authors:** Sanasar G. Babajanyan, Sofya K. Garushyants, Yuri I. Wolf, Eugene V. Koonin

## Abstract

Prokaryotes can acquire antivirus immunity via two fundamentally distinct types of processes: direct interaction with the virus as in CRISPR-Cas adaptive immunity systems and horizontal gene transfer (HGT) which is the main route of transmission of innate immunity systems. These routes of defense evolution are not mutually exclusive and can operate simultaneously, but empirical observations suggest that at least in some bacterial and archaeal species, one or the other route dominates the defense landscape. We hypothesized that the observed dichotomy stems from different life-history tradeoffs characteristic of these organisms. To test this hypothesis, we analyzed a mathematical model of a well-mixed prokaryote population under a stochastically changing viral prevalence. Optimization of the long-term population growth rate reveals two contrasting modes of defense evolution. In stable, predictable and fluctuating, unpredictable environments with a moderate viral prevalence, direct interaction with the virus and horizontal transfer of defense genes become the optimal routes of immunity acquisition, respectively. In the HGT-dominant mode, we observed a universal distribution of the fraction of microbes with different immune repertoires. Under very low virus prevalence, the cost of immunity exceeds the benefits such that the optimal state of a prokaryote is complete defense systems. By contrast, under very high virus prevalence, horizontal spread of defense systems dominates regardless of the stability of the virome. These findings might explain consistent but enigmatic patterns in the spread of antivirus defense systems among prokaryotes such as the ubiquity of adaptive immunity in hyperthermophiles contrasting their patchy distribution among mesophiles.

**IMPORTANCE:** The virus-host arms race is a major component of the evolutionary process in all organisms that drove the evolution of a broad variety of immune mechanisms. In the last few years, over 200 distinct antivirus defense systems have been discovered in prokaryotes. There are two major modes of immunity acquisition: innate immune systems spread through microbial populations via horizontal gene transfer (HGT) whereas adaptive-type immune systems acquire immunity via direct interaction with the virus. We developed a mathematical model to explore the short term evolution of prokaryotic immunity and show that in stable environments with predictable viral repertoires, adaptive-type immunity is the optimal defense strategy whereas in fluctuating environments with unpredictable virus composition, HGT dominates the immune landscape.

## INTRODUCTION

Bacteriophages (phages for short), viruses infecting bacteria, are the most abundant biological entities on Earth (1–4). Phages are present in all habitats, and in many of these, the number of virus particles exceed that of bacterial cells several-fold (4–8). Phages have been estimated to cause about 20% of the daily bacterial mortality in the ocean (9). Thus, antivirus defense is central for the survival and evolutionary success of bacteria as well as archaea. Indeed, through the 4 billion years of incessant virus-host arms race, prokaryotes have evolved an enormous variety of defense mechanisms. The recent development of computational and experimental pipelines for identification of defense systems in prokaryotes led to the discovery of more than 200 distinct defense systems, and additional ones are being found at a high pace (10, 11){Millman, 2022 #1895).

Despite the striking overall diversity of the antiphage defense systems found in different bacteria, each individual genome carries, on average, only 7-9 identifiable ones (12). The limited number of defense systems per genome is generally attributed to the fitness cost associated with antivirus immunity, on one hand, and the population-level defense provided by the diverse immune repertoire, on the other hand (13). Short term evolutionary experiments reveal a complex picture whereby some defense systems are costly whereas others do not appear to affect bacterial cell fitness in the absence of phage infection (14). Nevertheless, apart from the generic cost of replication and constitutive (even if low level) expression of defense systems, autoimmunity can be a substantial fitness burden as demonstrated, for example, in competition experiments with bacterial strains carrying or lacking a particular restriction-modification (RM) system (15). Bacteria carrying other defense systems, for example Lit, also have been shown to growth disadvantage on bacteria in the absence of phage (16). Furthermore, CRISPR systems have been shown to erroneously introduce self-targeting spacers at substantial rates, resulting in autoimmunity (17–22). More generally, frequent loss of various defense systems in bacterial evolution implies that antiphage immunity is a major fitness burden for prokaryotes (23). Even closely related strains of bacteria of the same species often carry different sets of defense systems, pointing to a high turnover rate of the prokaryotic defensome (14, 24–26). Together, these observations prompted the development of the pan-immune system concept according to which, although individual strains cannot carry all or most of the defense systems because of the fitness burden that would outweigh the benefits of immunity, they continuously vary their immune repertoires by acquire defense genes from closely related strains via horizontal gene transfer (HGT) (27).

Prokaryotic defense systems show enormous variability of the mechanisms of action and employ all types of cellular machinery to protect microbial populations from viruses (12, 28). On the most basic level, defense mechanisms can be classified into innate and adaptive immunity. Innate immunity mechanisms prevent virus reproduction in infected cells or induce dormancy or death of infected cells via a broad variety of molecular mechanisms. The most common type of innate immunity are the RM systems that recognize and destroy unmodified viral DNA, while protecting the host DNA through specific base modification, or conversely, by recognizing and cleaving viral DNA with modified bases. In the last few years, systematic efforts using dedicated computational and experimental pipelines have led to the discovery of numerous innate immunity systems that widely differ in their specificity, often providing protection only against a narrow range of viruses (12, 28). The fundamental unifying feature of innate immunity mechanisms is that they are pre-adapted to block reproduction of a certain range of viruses (or in some case, plasmids) and do not adjust their specificity in response to the infection by a particular virus.

The second major type of antivirus defense is adaptive immunity whereby the bacterial genome undergoes a specific modification after an encounter with a virus, and as a result, the cell and its descendants become immune against subsequent infections with the given virus and possibly its close relatives. The only known form of adaptive immunity in prokaryotes is represented by the widespread CRISPR-Cas systems (hereafter CRISPR for brevity) (29–32). When a viral or plasmid DNA penetrates a cell encoding a CRISPR system, a dedicated complex of Cas proteins excises short DNA fragments from invading viruses or plasmids and inserts them into the CRISPR loci between the repeats (these unique inserts are hence known as spacers). If the cell survives infection, for example, being infected by a defective phage, processed CRISPR locus transcripts function as guide RNAs protecting the cell and its progeny from the cognate virus by targeting Cas nucleases to the invading DNA. Although so far CRISPR remain the only experimentally characterized adaptive immune systems in prokaryotes, the recent discovery of TIGR-Tas systems that are unrelated to CRISPR but are also capable of RNA-guided target cleavage (33) suggests that additional adaptive immunity mechanisms remain to be discovered.

A fundamental distinction between innate immunity and adaptive immunity is the difference between the modes of immunity acquisition. While adaptive immunity is acquired through direct interaction with the invading pathogens (their nucleic acids, in the case of CRISPR system), innate immunity systems spread among prokaryotes via HGT which is the defining process in the evolution of prokaryotes and is often mediated by mobile genetic elements (MGE) including plasmids and viruses (34, 35). High horizontal mobility indeed has been demonstrated for defense-related genes compared to other functional classes of genes in prokaryotes (24–26). Gain of immunity via direct interaction with a virus might not be limited to adaptive immunity proper. For example, resistance-conferring mutations in cell surface proteins that play a major role in bacterial defense against phages and can be mildly deleterious in the absence of the virus (36, 37) would sweep the population once a virus wipes out the susceptible cells. This is a route of immunity acquisition that does not involve HGT, but results from interaction with the virus and in this respect resembles adaptive rather than innate immunity acquisition.

The abundance of prokaryotic defense systems varies widely across habitats, and as could have been expected, bacteria from environments with higher virus load tend to carry more defense systems in their genomes than those from virus-poor environments (38). There are also distinct patterns of distribution of different types of defense systems across prokaryotes with different lifestyles. The most prominent of these seems to be the high prevalence of CRISPR in bacterial and archaeal thermophiles, including near ubiquity among hyperthermophiles, in contrast to mesophiles among which less than a half possess CRISPR (32).

Analysis of agent-based mathematical models accounting for the fitness cost of the adaptive immunity suggested that adaptive immunity (CRISPR) is highly beneficial at intermediate viral diversity, which seems to be the case in hyperthermal habitats, whereas at both low and high diversity, innate immunity was predicted to be the dominant form of defense (39, 40). Recently, analysis of a mathematical model factoring in metabolic and autoimmune costs of different bacterial defense systems within a generic Lotka-Volterra population dynamics framework yielded an upper boundary of the optimal number of defense systems per genome that appears to be compatible with genome analysis results (41).

Notwithstanding the work mentioned above, general understanding of the ecological factors defining the distributions of adaptive and innate immunity systems in prokaryotes is lacking. We sought to address this problem by performing a cost-benefit analysis of the two major modes of immunity acquisition in prokaryotes, direct interaction with a virus (DIV) and HGT, depending on the virus prevalence and the temporal stability of the local virome. Analysis of our general mathematical model of virus-host coevolution shows that DIV is beneficial in temporally stable, predictable environments with moderate virus prevalence. At very low virus prevalence, the optimal state of a prokaryotic population is complete absence of defense systems, whereas at very high virus prevalence, HGT dominates.

## RESULTS

### The model of immunity acquisition in prokaryotes

We consider the dynamics of a well-mixed population of microbes under time-dependent virome composition which fluctuates due to the random appearance and disappearance of viruses in the environment, with fixed virus pool size *L*. Each virus is described by the probability of its presence at time *t* (μ_*α*_), viral prevalence, and its characteristic time of viral dynamics *τ*_*α*_, where *α* = 1, ‥, *L*, respectively.

These parameters are determined by the appearance and disappearance probabilities of the given virus *α, q*_*α*_ and *p*_*α*_, respectively, through the following relations

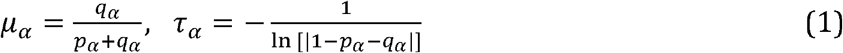

where *μ*_*α*_ is the probability of the presence of the virus in the stationary state, and *τ*_*α*_ is the characteristic time that reflects the predictability of the dynamics of the given virus. If *p*_*α*,_ *q*_*α*_ ≪1 the current state of the virus, that is, presence or absence, is likely to persist in the subsequent generations. Conversely, if *p*_*α*,_ *q*_*α*_ ∼ 1 then, the current state of the virus will likely change every generation (from presence to absence and vice versa). In both cases, the dynamics of the virus is predictable for a longer time than it is when *p*_*α+*_ *q*_*α*_ ∼ 1. In the latter case, the pathogen dynamics quickly converges to the stationary state *μ*_*α*_, that is, the correlation between the states of the virus across generations declines fast. Thus, if all viruses have long or short characteristic times, the state of the virome is, respectively, more or less predictable (see section I in SI Appendix for further details).

Within the scope of the model, the routes of antivirus immunity acquisition by prokaryotes belong to two classes. The first class includes a broad range of relatively complex innate immunity systems that cannot emerge *de novo* in any given population. These defense systems evolved over billions of years and disseminate across the bacterial and archaeal diversity via horizontal gene transfer (HGT). Within any population, the dynamics of such immune systems is Wrightian (42), that is, dominated by the balance between the rates of gene gain and loss which are neutral processes independent of the state of the environment (effectively, the virome composition, in the model) and the environment-dependent differential survival of carriers and non-carriers of a particular system (a selective phenomenon). The second class covers acquisition of defense resulting from a more or less direct interaction with the virus (DIV), either via adaptive immunity mechanisms, as in the case of CRISPR systems, or via a mutation process (broadly defined) whereby mutations can occur but are highly unlikely to persist in the population even for a short time in the absence of the cognate virus, as in the case of cell surface proteins recognized by viruses. In these cases, the evolution is quasi-Lamarckian (42), that is, defense emerges as adaptation in response to the virus presence in the environment and is inherited by the progeny (even in the absence of the virus) until lost by chance.

To simplify the model framework, we represent the immune status of the organism as a genotype vector, that is, the immune repertoire that indicates immunity against a given virus without distinguishing between the routes of defense acquisition (HGT or DIV). The immune repertoire of a cell is a binary vector of length *L*, ***s***, that represents whether the cell is protected against each of the *L* viruses or not. A cell is immune to virus *α* if *s*_*α*_ = 1 and susceptible if *s*_*α*_ = 0 in ***s***, where *α* = 1, ‥,*L*,. The cells, however, can independently employ either HGT or DIV, or both to acquire immunity against a virus. Thus, we assume that the defenses acquired through either HGT or DIV are indistinguishable within the model, that is, behave in the same way, regardless of the route of their acquisition, thus avoiding complications caused by the separation of defense genes acquired via the two distinct routes‥

Through the defense acquisition processes, HGT and DIV, cells with different immune repertoires appear over time which, in the model, is the only difference between cells in the population. The fractions of cells with different immune repertoires vary over time due to the competition for common resources and acquisition, and loss of defense genes. It is assumed that acquisition and loss of defense genes do not depend on the current immune repertoire of the cell and the probability of losing a defense gene *π*_01_ is fixed, being independent of both the environment and population composition.

The probability of acquiring immunity against any virus depends both on the current state of the virus prevalence in the environment and the population composition (distribution of immune repertoires) through the two distinct mechanisms of immunity acquisition, DIV and HGT. Any cell that is currently not immune to a virus *α*, which is present in the environment, can become immune to it via DIV with probability *p*_*DIV*_. An alternative mechanism for defense acquisition is through HGT, which is assumed to operate as a copy-paste mechanism, that is, without gene loss. The acquisition of immunity through HGT is characterized by *p*_*HGT*_, the conditional probability that the cell acquires immunity to the virus *α* once a donor, that is, a cell with *s*_*α*_ = 1, is encountered. The probability of encountering such a donor cell varies over time due to the change in the population composition. In contrast to defense acquisition via DIV, HGT allows a cell to acquire immunity to a virus that is not currently present in the environment.

Throughout this work, we assume that the host population is initially in a naive state. Technically, this means that all cells lack any defense genes. However, a non-zero probability *p*_*DIV*_, even if infinitesimally small, ensures the capacity of the cells to acquire immunity against any virus appearing in the population. These cells can serve as donors for spreading defense genes via HGT, given that the model does not differentiate between immunity acquired via DIV or HGT, that is, once acquired via either HGT or DIV immunity can be transmitted further.

Alternatively, it would be possible to assume that initially all defense genes are present in the environment, for example, assigning uniform fractions of cells with all possible immune repertoire, so that defense acquisition is still possible due to HGT, even if *p*_*DIV*_ = 0. To ensure that all simulation runs started from the same initial conditions, we elected to make the initial state of the population formally naive, but with non-zero *p*_*DIV*_, allowing immunity to emerge locally and spread via HGT.

The fitness of a cell in the environment ***e***(*t*) is defined as the product of the probabilities of reproduction and cell survival. The survival probability of a cell is affected by the virus prevalence in the environment. We assume that each virus present in the environment (*e*_*α*_ = 1) can kill a cell that is not immune to this particular virus (*s*_*α*_ = 0, where *α* = 1, ‥,*L*,), with probability *d*_*α*_ ∈ [0,1]. If a cell is immune to the given virus *α*, that is, *s*_*α*_ = 1, then, the presence of the virus in the environment does not decrease the survival probability of that cell. It is further assumed that each cell in the population may die with the probability *d*_0_, regardless of its immune repertoire and the presence of any virus. Immunity incurs a cost on the fitness of the cell by reduces the probability of cell reproduction. That is, each defense gene in the cell immune repertoire decreases the probability of cell reproduction by a factor 1 − *c*_*α*_, where *c*_*α*_ is the cost of immunity to the virus *α*(*c*_*α*_ ∈ [0,1]).

In addition to the cost of all defense genes, maintaining a mechanism for defense acquisition through DIV is assumed to come with an additional cost on cell reproduction. This cost is proportional to the efficiency of defense acquisition and is given by 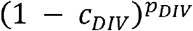 which is fixed for all cells in the population, regardless of their current immune repertoires. That is, all cells in the population possess the same mechanism of defense acquisition through DIV with a fixed efficiency. The last assumption is applicable particularly for adaptive immunity systems, such as CRISPR-Cas systems. It should be noted that defense acquisition via HGT can indirectly decrease the reproduction probability of a cell because the cell can acquire costly defense genes against viruses that are not present in the environment at the time of acquisition.

Our analysis of the model is based on finding the maximum long-term growth rate of the population by maximizing it with respect to (*p*_*HGT*,_*p*_*DIV*_), which are assumed to be population level traits, for the given virome composition (*μ*_*α*_, *τ*_*α*_), *α* = 1,…, *L*,.

### Optimal modes of defense acquisition

Stochastic fluctuations of the virome, that is, random appearance and disappearance of viruses in the environment, cause variation in cell fitness due to the changes in the cell survival probabilities caused by the changing virome composition. Considering the expected fitness of cells with immune repertoires ***s*** and ***s***^⍰^ that differ only with respect to virus *α*, that is, *s*_*α*_ = 0 and 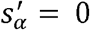, acquisition of defense against virus *α* is beneficial if

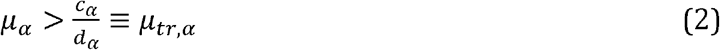

where μ_*α*_ is the probability of the presence of virus *α* in the environment at any time, *c*_*α*_ is the cost of defense against virus *α, d*_*α*_ is the probability of death caused by the virus (see section II in SI Appendix for details) and *μ*_*tr,α*_ is the threshold prevalence for this virus. Indeed, if the cell is immune or susceptible to the virus *α*, then, the average fitness reduction is 1 − *c*_*α*_ and 1 − *μ*_*α*_ *d*_*α*_, respectively. Thus, immunity against the given virus is beneficial when the decrease in average fitness due to the cost of defense is smaller than the fitness decrease caused by the lack of defense against the given virus. The ratio of the defense cost and lethality of a virus defines the threshold value for the probability of the virus presence above which acquiring defense against this virus becomes beneficial to the cell in terms of the average fitness. The maximum average fitness depends on the viral prevalence in the environment (Fig. 1A). If the parameters are the same for all viruses, that is, *c*_*α*_ = *c, d*_*α*_ = *d* and *μ*_*α*_ = *μ*, cells immune to all viruses have the highest average fitness when the probability of the presence of the viruses is above the threshold (2), whereas when this probability is below the threshold, naive cells have the highest average fitness. However, the optimal long-term growth of the population generally cannot be inferred from the average fitness of the cells because average fitness is defined only by the virus presence probabilities *μ*_*α*_ and is independent of the characteristic times *τ*_*α*_. The optimal mode of defense acquisition, however, depends on both these quantities. In environments with longer characteristic times *τ*_*α*_, where the environment-specific virome is more predictable, it might be optimal to adapt specifically to the current virome composition, rather than adapting to the average state, which is the optimal strategy if the environment switches frequently, that is, viruses appear and disappear often, as further discussed below.

**FIG. 1.**
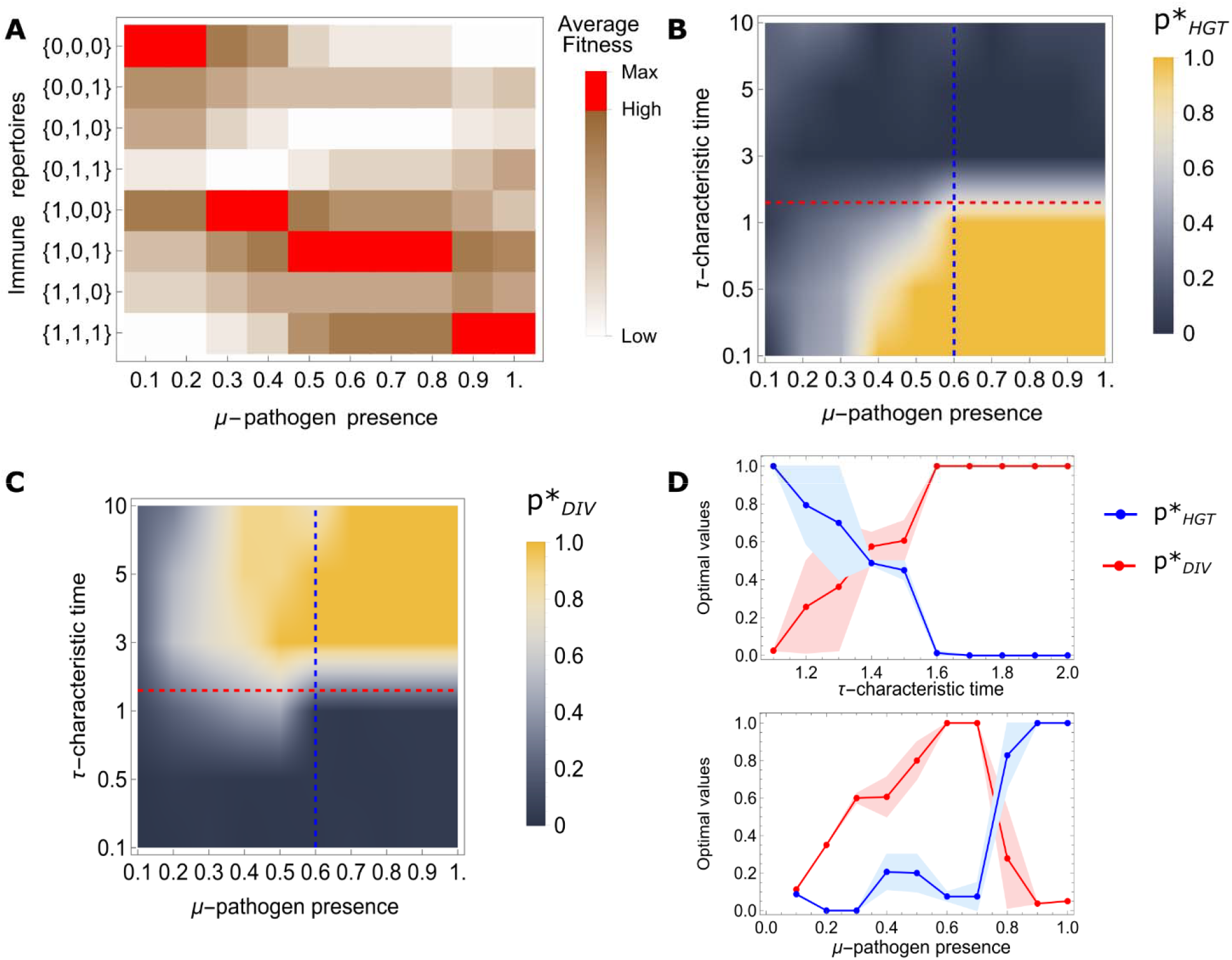
Phase space of optimal modes of defense acquisition depending on the probability of virus presence in the environment and characteristic time of viral dynamics. A) Ordering of the average fitness values of all possible immune repertoires against three viruses over virome composition fluctuations depending on the viral prevalence. Red squares denote the fittest immune repertoire in the given environment. All three viruses are assumed to have the same characteristic times, but different presence probabilities, respectively. These probabilities are constrained within the interval. The remaining model parameters are,,,. The threshold value of the presence (2) is the same for all viruses,. The population dynamics was followed for generations. B) Optimal values of the probability of defense acquisition via HGT depending on viral prevalence in the environment and characteristic time of viral dynamics C) Optimal values of the probability of defense acquisition via DIV depending on viral prevalence in the environment and characteristic time of viral dynamics D) Upper panel, crossing from the HGT-dominated regime to the DIV-dominated regime with increasing characteristic time of viral dynamics *τ*, μ = 0.6 (blue dashed line in B) and C)). Lower panel, crossing from the DIV-dominated regime to the HGT-dominated regime with increasing viral prevalence, *π* = 1.5 (along red dashed line in B) and C)).

For low viral prevalence in the environment, *μ*_*α*_ < *μ*tr, the optimal values of defense acquisition probabilities via both HGT and DIV are small (Fig.1B and Fig.1C). In the extreme case of a very low viral prevalence, defense acquisition is deleterious, so that 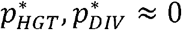 and the naïve population has the highest long-term fitness. Increasing virus prevalence leads to either the HGT-dominated or the DIV-dominated regimes of defense acquisition, depending on the characteristic times of viral dynamics. In frequently fluctuating, unpredictable environments (short characteristic times), the optimal route of defense acquisition is HGT, whereas in more predictable environments with longer characteristic times, DIV becomes the optimal defense strategy (Fig.1B,C). In extreme cases, the optimal values are close to 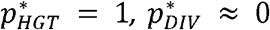 and 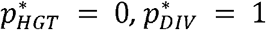 for frequently fluctuating and highly stable environments, respectively.

The HGT-dominated and DIV-dominated regions of the phase space are separated by a narrow range of values with intermediate environment fluctuation frequency in which both modes of defense acquisition operate concomitantly. With increasing characteristic times of viruses, the environment becomes more predictable, and at moderate viral prevalence, there is a sharp transition from the HGT-dominated to the DIV-dominated regime (upper panel in Fig.1D). We further examined the dependence of the optimal values of defense acquisition probabilities via HGT and via DIV on the viral prevalence in the environment for moderate values of the characteristic times of virus dynamics (lower panel in Fig.1D), that is, in the region between the areas dominated by the two modes of defense acquisition. In this narrow “valley” of the parameter space, where both modes operate, the outcome of stochastic optimization of the long-term growth of the population becomes unstable so that the outcomes differ between simulations (in Fig. 1, we present the mean of the outputs of two runs with respective deviations).

At long characteristic times of viral dynamics, once the probability of presence of at least one virus exceeds the threshold value (2), DIV becomes the dominant mode of defense acquisition. Long characteristic time means that the virome composition is predictable, that is, the currently present viruses typically persist for many generations, hence, DIV (in particular, adaptive immunity) allows the cells to acquire immunity specifically against the present viruses. Thus, the cells with immune repertoires that match the current virus diversity become more prevalent in the population. By contrast, if defense is acquired stochastically, via HGT, cells with non-matching immune repertoires will appear, decreasing the long-term growth rate of the population. Therefore, in this regime, DIV is the optimal strategy of defense acquisition in spite of the associated cost 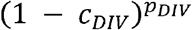. Clearly, the maintenance cost of adaptive immunity affects the range of optimal values of *p*_*DIV*_. That is, increasing cost *c*_*DIV*_ decreases the range of possible values of *p*_*DIV*_ that provide fitness advantage over the naïve population or HGT-dominated strategies. If the additional cost of DIV immunity acquisition is higher than the threshold, it might be beneficial for the population to dispense of adaptive immunity and instead stay in the naïve state. Conversely, the range of *p*_*DIV*_ values that provide long-term fitness advantage compared to the naïve population increases with the number of viruses and viral prevalence of the environment (see section IV in SI Appendix for further details).

By contrast, in unpredictable environments with short characteristic times of viral dynamics, paying the extra cost of the DIV acquisition mode is not advantageous because the environment is unlikely to persist for many generations, so that cells with immune repertoires matching the current environment will be at a disadvantage in the next generations. Thus, it is preferable to acquire immunity via HGT, avoiding the cost of maintenance of adaptive immunity. Furthermore, in frequently changing environments, if immunity spreads via HGT, cells that are at a disadvantage due to a suboptimal immune repertoire, can become the fittest ones in the next generations as the virome changes.

The optimal values of the rates of the two processes of immunity acquisition, 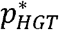 and 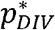, also depend on the gene loss probability *π* _01_. For small gene loss probabilities, the optimal immunity acquisition rates are smaller than for high loss probabilities (see SI Figure 4). Indeed, high values of and ensure the emergence of the optimal immune repertoire of the population in the current environment. However, once the environment changes and the optimum immune repertoire changes with it, the lower gene loss probability will delay the adjustment. Thus, the optimal values of defense acquisition probabilities and need to be in balance with the gene loss probability.

In the case of three viruses with different presence probabilities considered in Fig. 1, the second virus, with, appears rarely, even in a high viral prevalence () environment, where the remaining viruses are almost always present. Nevertheless, DIV provides hosts with the flexibility to adapt even to this comparatively rarely present virus such that it is not necessary to possess immunity against all viruses simultaneously (see Fig.2A) although the average fitness of the fully immune cells is the highest (Fig. 1A). Thus, the extended periods of presence and absence of this virus provide for DIV being the optimal defense acquisition mode despite the extra cost of its maintenance. If HGT operates in this case, too, there will be greater fractions of cells that are immune against the second virus in the periods when it is absent in the environment.

**FIG. 2.**
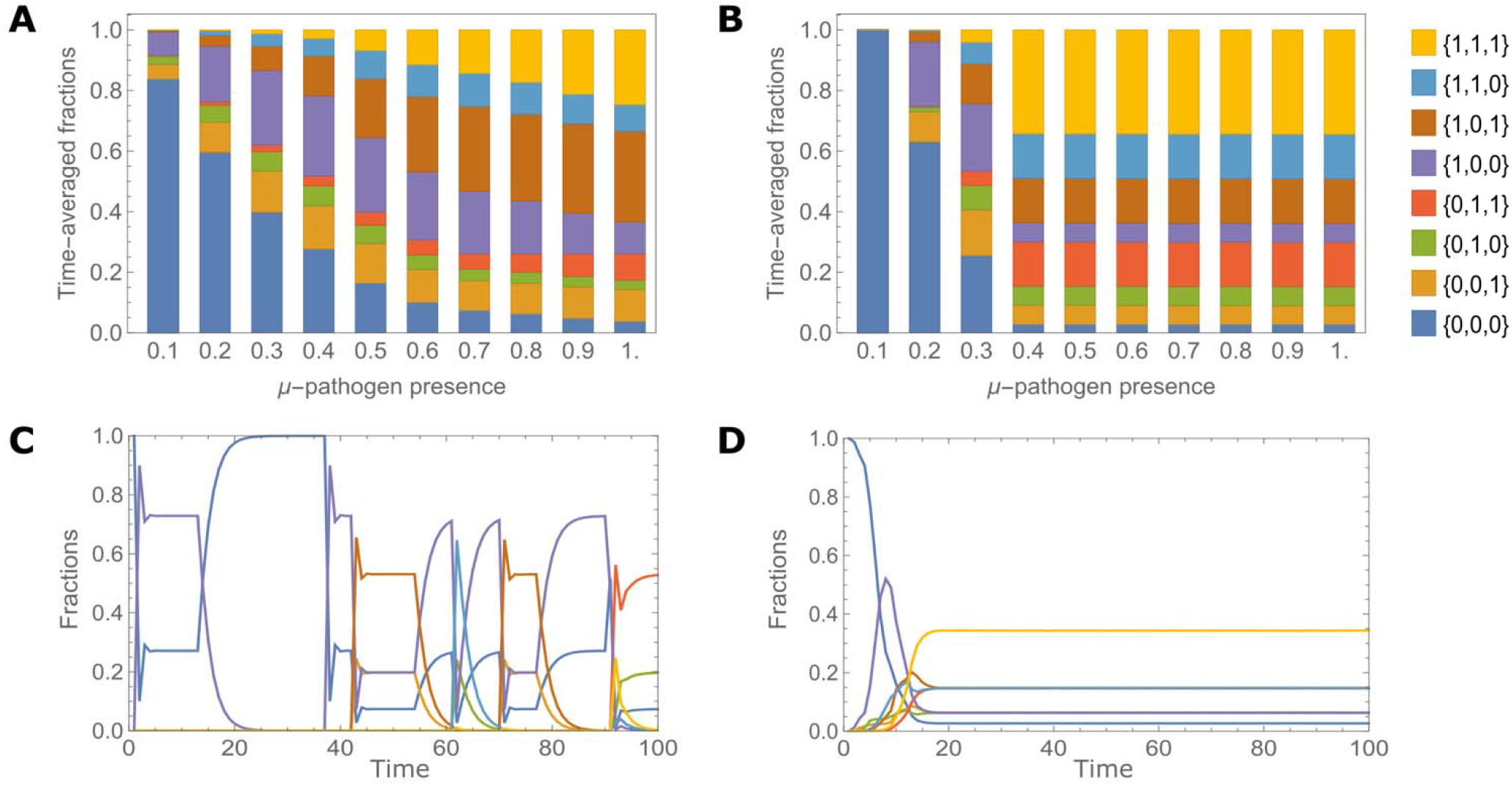
Distributions of immune repertoires in the population corresponding to different modes of optimal defense acquisition process depending on the viral prevalence in the environment. A) Composition of the population, that is fractions of cells with different immune repertoires averaged over the time *T* for a long characteristic time of viruses, *τ*_*α*_ = 10, *α* = 1,2,3. The optimal values of defense acquisition probabilities are taken from Fig.1B and Fig.1C. Increasing viral prevalence in the environment with long characteristic time leads to DIV-dominant immunity, see Fig.1C. B) Composition of the population of cells for a short characteristic time of viruses, *τ*_*α*_ = 0, *α* = 1,2,3. Increasing viral prevalence in the environment with short characteristic time leads to HGT-dominated region of defense acquisition, see Fig.1B. C) Time-dependence of the fractions of cells with different immune repertoires for a long characteristic time, *τ*_*α*_ = 10, *μ* = 0.5. D) Time-dependence of the fractions of cells with different immune repertoires for a short characteristic time of viruses, *τ*_*α*_ = 10, *μ* = 0.5. The colored areas (see the key at B) in all panels represent fraction of cells with the given immune repertoire. The remaining model parameters are the same as in Fig. 1. The trajectories shown in C (and D) were generated for the optimal values of defense acquisition probabilities through HGT and DIV. The population dynamics was analyzed for *T* = 10^4^ generations; in C) and D), the population dynamics is shown for 100 generations.

If the probabilities of presence are high for all viruses, HGT becomes the dominant mode of defense acquisition even in predictable environments with long characteristic times of viral dynamics (SI Figure 2). Indeed, when the presence probabilities of all viruses are high,, the environment becomes effectively fixed, with all viruses present. Under these conditions, acquisition of immunity via HGT instead of DIV will be favored due to the absence of the extra maintenance cost.

If all viruses have different characteristic times and presence probabilities, the phase space of optimal values of HGT and adaptive immunity becomes more complex. Consider again the case of different presence probabilities for three viruses {*μ*_1_, *μ*_2_, *μ*_3_} = {3/2*μ, μ*/2, *μ*}, now with the characteristic time of the least frequent virus fixed at *τ*_2_ = 1. We set the characteristic times of the two remaining viruses either as *τ*_3_ > *τ*_2_ > *τ*_1_ or as *τ*_1_ > *τ*_2_ > *τ*_3_ (SI Figure 3).

In the former case, the virus with the highest probability of presence (*μ*_1_ ∼ 1) has the shortest characteristic time (*τ*_1_ < 1), whereas the second most frequent virus has a longer characteristic time (*τ*_3_ > 1). If the probability of presence of the least frequent virus is below the threshold value (*μ*_2_ = /2< *μtr*), then, in this environment, DIV becomes the optimal mode of defense acquisition because acquisition of defense against the infrequently present virus, the second pathogen, is disadvantageous whereas it still allows to be immune against the most prevalent and the most predictable pathogens, the first and third pathogens, respectively. Indeed, the presence of a predictable virus with above threshold prevalence, that is, the third virus, ensures the optimality of DIV mode because there will be longer presence and absence periods of that virus. Increasing the frequency of the least frequent virus over the threshold value (*μ*_2_ = *μ*/2 > *μ*_*tr*_), hence further increasing the prevalence of all pathogens, switches the optimal mode of defense acquisition to HGT, avoiding the extra cost of the DIV capability. In the latter case (*τ*_1_ > 1, *τ*_3_ < 1), where the most prevalent virus is also the most predictable, that is, is expected to be present over long time periods, whereas the second most prevalent virus is the least predictable, DIV is no longer beneficial, and HGT acquisition, again, becomes optimal. Thus, DIV allows fine tuning to more complex environments where it is not beneficial to acquire defense genes against rarely present viruses, but there are still viruses that are more predictable even if not highly prevalent.

When all the viruses have short characteristic times, even with moderate values of the probabilities of virus presence, the virus dynamics quickly converges to the stationary state. Thus, the population dynamics is solely determined by mean fitness values of different immune repertoires. The optimal mode of defense acquisition under these conditions is 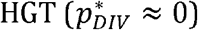, and acquisition of immunity against a given becomes decoupled from the instantaneous state of the environment. In this regime the probabilities of defense acquisition against any pathogen are only determined by the population composition that is driven by the selection process, where the fitness of cells is determined by their expected values over the fluctuation of virome composition (see section VI in SI Appendix).

In the HGT-dominated mode, the composition of the population at the optimal defense acquisition probabilities (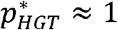 and 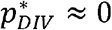) converges to the equilibrium where the fractions of cells with different immune repertoires are given by

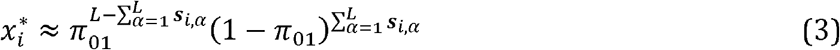

Here, *x*_*i*_ is the fraction of cells with immune repertoire ***s***_*i*_, *i* = 1,…,2^*L*^ *π*_01_ is the gene loss probability, which is the same for all genes, and 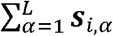 is the number of defense genes in the immune repertoire ***s***_*i*_. The equilibrium (3) arises as long as the population dynamics is described by eq. (8) in SI (see SI VI for further details), where the fitness values of cells are replaced by the respective expectations. At equilibrium (3), the probability of acquiring any defense gene becomes equal to 1 − *π*_01_ due to the composition of the population (see Section VII in SI), that is, complementary to loss probabilities. Thus, in the unpredictable environment, the composition of the population tends to (3), where the acquisition probability of any defense gene is determined only by the gene loss probability, *π*_01_, and the size of the virus pool, *L*. The most prevalent cells at the equilibrium possess *L*(1 − *π*_01_) different defense genes on average. When viruses have identical parameters (μ_*α*_ = μ, *c*_*α*_ = *c* and *d*_*α*_ = *d*), the long-term fitness of the population at equilibrium has the following simple form (see Section VII in SI)

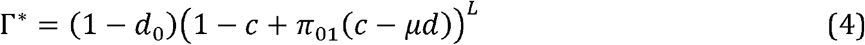

Here *c*≤ μ*d*, otherwise, acquisition of any defense genes is deleterious, hence equilibrium (3), is not optimal anymore and cannot be reached. In an unpredictable environment (short characteristic time), the time-averaged immune landscape of the population is dominated by defenseless cells at low virus prevalence, but once the threshold (2) is reached, that is, in the HGT-dominated regime, the fractions of cells with different immune repertoires (3) reach a stable equilibrium independent of μ (Fig. 2B,D).

Conversely, in the DIV-dominated regime, the immunity landscape of the population changes over a much wider range of viral prevalences (Fig. 2A, C). In this case, the distribution of the immune repertoires is not determined only by the average fitness, but rather, critically depends on the characteristic times of viruses. Furthermore, in this mode, the immune repertoire associated with the highest average fitness is not necessarily the most prevalent one (see the distributions for μ = 0.9 and μ = 1 in Fig. 2A).

## DISCUSSION

In this work, we develop a virus-host coevolution model to analyze the interplay between HGT of immune systems within a microbial population and acquisition of immunity via direct interaction with viruses. We show that the choice of the optimal defense strategy depends, primarily, on the diversity and temporal stability of the virome, as well as the rates of gene gain and loss by the prokaryotic hosts (Figure 3). In environments with a moderate viral prevalence and predictable virome behavior, acquisition of defense via direct interaction with the viruses, in particular adaptive immunity, becomes the optimal strategy. In less predictable, fast fluctuating environments or under very high virus prevalence, the defense landscape is dominated by immune systems that are acquired via HGT. At low virus prevalence, the optimal strategy is having no defense systems at all. This appears unexpected but, arguably, in the context of population-level selection, it is preferable to let a large fraction of the population die due to infrequent viral infection, with the remaining cells leading the rebound once the virus clears, rather than pay the fitness cost of defense in the absence of viruses. At high virus prevalence, even in slowly fluctuating, predictable environments, HGT of immunity systems dominates over the defense landscape, apparently, because, after exceeding a threshold value, diverse viruses overwhelm adaptive immunity. The modeling results presented here are generally compatible with those of a previous study which concluded that in prokaryotes, adaptive immunity (CRISPR) should be beneficial within a mid-range of virus diversity in the environment (39). Compared to that early work, here we explore the complete phase space of the virus-host system showing how the preferred mode of immunity depends on the prevalence of multiple viruses and temporal stability of the virome.

**FIG. 3.**
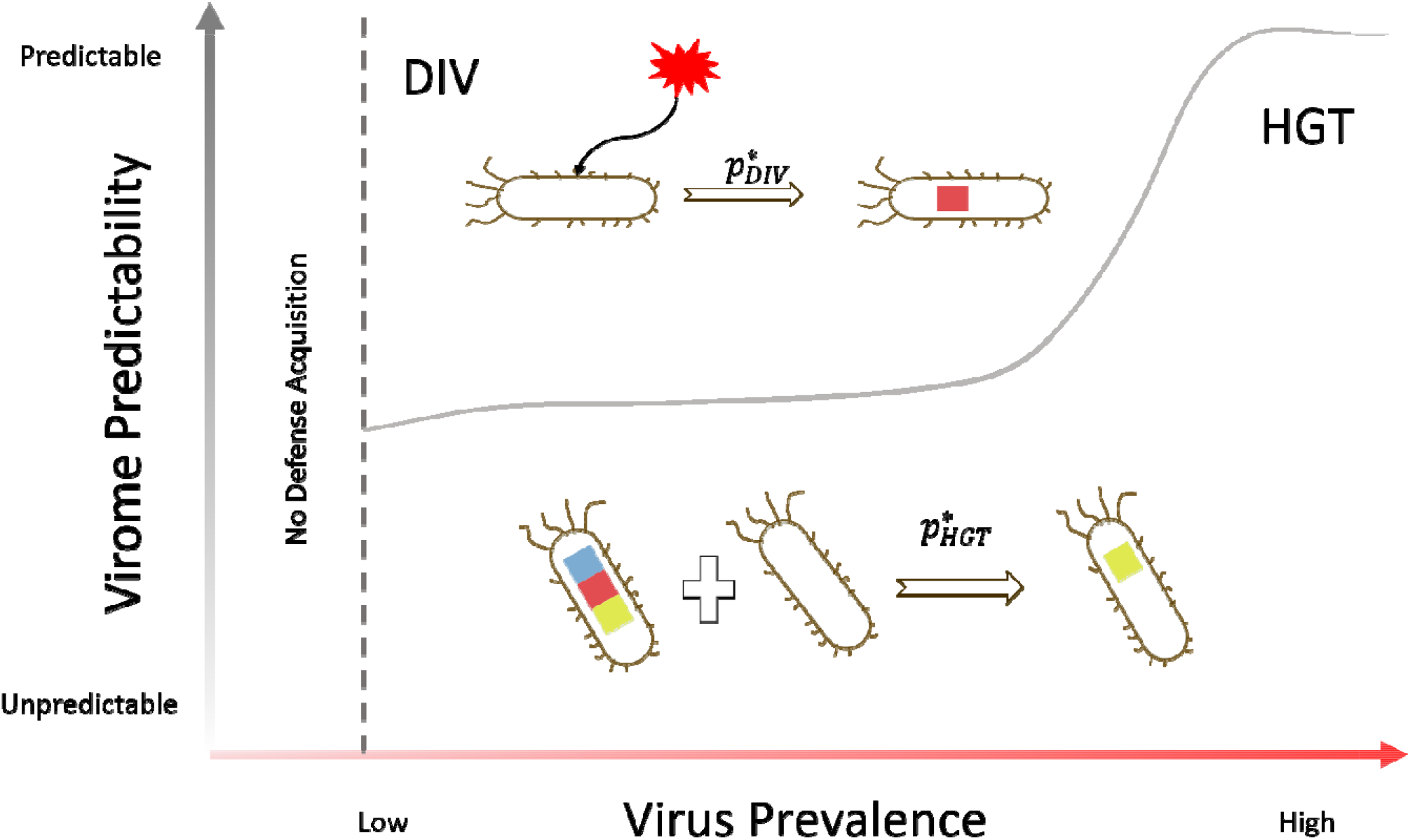
Optimal modes of defense acquisition depending on virus prevalence and virome stability. The schematic shows that, in environments with a moderate viral prevalence and predictable virome behavior, DIV, in particular adaptive immunity, is the optimal strategy. In less predictable, fast fluctuating environments and/or at very high virus prevalence, the defense landscape is dominated by immune systems that are acquired via HGT. At low virus prevalence, the optimal strategy is having no defense systems at all.

In fast fluctuating environments, where the defense landscape is dominated by HGT of immune systems, the distribution of the fractions of cells with different immune repertoires reaches equilibrium and remains constant once the viral prevalence exceeds a threshold value. This distribution depends only on the gross parameters of the system, namely, the total number of viruses and the probability of gene loss, but not on the fitness values of the cells with different immune repertoires and hence not on the cost of defense. At equilibrium, defense genes are transferred among the cells in the population, so that each cell can gain any defense gene with the same probability that is determined by the gene loss rate, implying a constant HGT rate of defense genes that provides the maximum attainable level of immunity at the population level.

The model presented here makes testable predictions regarding the absolute and relative prevalence of defense systems in prokaryotes in different habitats depending on viral prevalence, and the temporal stability of the virome. At present, quantitative information on these characteristics of different habitats is scarce, but qualitatively, the model seems to help explaining the most striking and still enigmatic features of the CRISPR adaptive immunity distribution across bacteria and archaea, namely, the near ubiquity of CRISPR among hyperthermophiles contrasting the patchy distribution among mesophiles (32). Indeed, hyperthermal environments match the conditions for the dominance of DIV including adaptive immunity that follow from our model, namely, a stable, predictable virome composition and moderate viral prevalence (43, 44).

The predictions of the model are also generally compatible with comparative genomics data. Specifically, analysis of the representation of different classes of defense systems across bacterial and archaeal phyla showed that enrichment in CRISPR typically was accompanied by depletion of innate immunity systems, such as RM, and vice versa (45). It should be pointed out, however, that whereas the model predicts that under most parameter combinations, either innate immunity disseminated via HGT or adaptive-like immunity acquired via DIV would be the sole mode of antivirus defense, with the mixed strategy existing only in a small region of the parameter space, comparative genomic data do not seem to match this prediction. More than half of the bacteria lack CRISPR(32) (even though they likely possess other forms of immunity acquisition via DIV), suggesting that they thrive in the parameter space region dominated by immunity acquisition via HGT, that is, unstable viromes and/or high virus prevalence. However, virtually none (with the exception of some intracellular symbionts and parasites) altogether lack any forms of immunity (12, 23). This discrepancy between the model predictions and observations implies either that the parameter combinations corresponding to the DIV-dominated regime of immunity are rare in natural environments or that the model fails to account for additional factors affecting the immune repertoire of prokaryotes. In the near future, detailed study of virome diversity and temporal dynamics in diverse habitats should allow quantitative testing of the predictions of the model.

## METHODS

### Population dynamics

The state of the environment is described by a binary vector of length *L*, that is, ***e***(*t*) = {*e*_1_(*t*), *e*_2_(*t*), *…, e*_*L*_ (*t*)}, where *L* is the pool size of the viruses. Each element in ***e***(*t*) indicates whether the virus *α* = 1,…, *L* is present at time *t* in the environment or not, *e*_*α*_ (*t*) = 1 and *e*_*α*_ (*t*) = 0, respectively. Each virus evolves independently of other viruses over time with probabilities of appearance and disappearance in the environment *q*_*α*_ and *p*_*α*_, respectively. Thus, the probability that the environment switches from ***e***(*t*) to ***e***′(*t+1*) is given by the following Expression 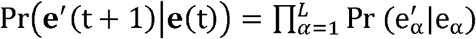, where each term in the product, depending on the values of 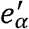 and *e*_*α*_, is equal 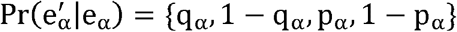.

In the environment of *L* viruses, there are 2^*L*^ possible different immune repertoires, ***S***_*i*_, where *i* = 1,…,2^*L*^. The fraction of cells carrying an immune repertoire ***S***_*i*_ at time *t* is denoted by *x*_*i*_(*t*), *i* = 1,…,2^*L*^. In what follows, we model the long-term behavior of the fractions of cells in the population carrying different immune repertoires, that is, the composition of population, represented by 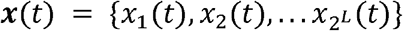. The composition of the population changes over time due to the defense acquisition, loss and competition for common resources.

We assume that acquisition and loss of defense genes are independent for each defense gene in the immune repertoire. The transition probability for a cell currently having immune repertoire ***S*** to any other immune repertoire, ***S***′, at time *t* is given by

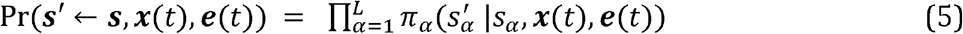

Where 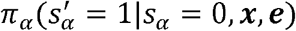 and 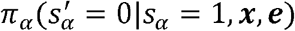 are the probabilities of acquiring and losing immunity to a virus *α* for the given environment, ***e***(*t*) and the composition of microbial population, ***x***(*t*), respectively.

By assumption, 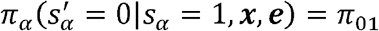, with 1− *π*_01_ being the probability of not losing immunity to a given virus.

The probability of defense acquisition against a virus *α*, either through adaptive immunity or horizontal transfer of the defense gene, is given by the following expression

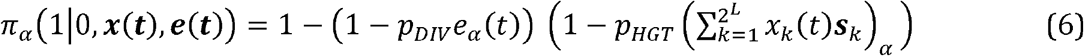

here, *p*_*DIV*_ ∈ [0,1] is the probability that a cell acquires immunity to a virus *α*, if the virus is present in the environment at the time *t, e*_*α*_ (*t*) = 1, that is, a direct interaction with the virus. Otherwise, if *e*_*α*_ (*t*) = 0, no defense acquisition occurs through direct interaction.

In (6), the sum is a vector of length *L*, where each term shows the fraction of the population that is immune to a particular virus *α, α* = 1,.., *L*. For each virus *α* there are 2^*L*−1^ different possible immune repertoires, ***s***, that are immune against it. That is *s*_*α*_ = *1* in all these immune repertoires. However, the fractions of cells that carry these repertoires vary over time. Thus, each term of the resulting vector shows the probability that a randomly selected cell carries immunity against a given virus *α, α* = 1,.., *L*, that is, the probability of choosing a cell that can serve as a donor for horizontal transfer of the defense gene against *α* = 1,.., *L* viruses. *p*_*HGT*_ shows the probability of successful horizontal gene transfer between recipient and donor cells, that is, the recipient cell acquires immunity to the virus *α*, once a donor for the HGT has been selected from the population.

The fitness of a cell is the product of reproduction and survival probabilities of the cell. For a cell with immune repertoire ***s*** it is equal to

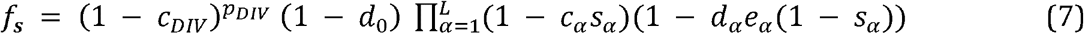

In (7), 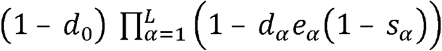 shows the survival probability of the cell with immune repertoire ***s*** in the environment ***e***(*t*). By assumption, a naive cell will reproduce with probability 1 (up to the fixed term of adaptive immunity cost), if survives, then the probability of reproduction of a cell with repertoire ***s*** is 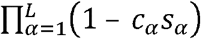. Here, *c* _*α*_ ∈ [0,1] is the cost of immunity against *α* virus, *s*_*α*_ = 1,0 shows wether the defense gene is present in the immune repertoire ***s*** or not. Note, that the reproduction probability of a given cell is independent on the environment ***e***, and is only defined by the immune repertoire of the cell, in contrast to survival probability that depends on the environment. Additionally, we assume that defense acquisition process through direct interaction with the viruses impose an additional cost 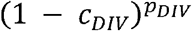, which is the same for all cells in the population.

We choose the multiplicative cost on the reproduction because the reproduction probability is always positive irrespective of the number of defense genes in the repertoire of the cell and their costs, in contrast to the additive cost 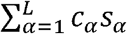. In this case, there will be an additional constraint on the cost *c*_*α*_ because the reproduction probability should be non-negative 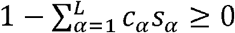. Additionally, the multiplicative cost resembles synergy between defense systems, since the reproduction probability of the cell is greater for multiplicative cost than for additive cost, that is any next defense gene has less overall impact on the reproduction for the case of multiplicative cost compared to additive cost of the defense genes (see section III SI for details).

Combining defense acquisition, loss and competition for common resources, we can write down the expression for the variation of the fractions of cells carrying all possible immune repertoires in the environment ***e***(*t*). These variations are given by the following dynamical system

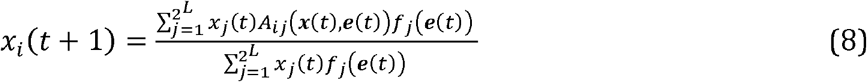

here *A*_*ij*_(***x***(*t*), ***e***(*t*)) ≥

0 is the transition probability of ***s***_*i*_ → ***s***_*j*_, where *i,j* = 1,…,2*L*, and 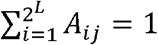. Note, that 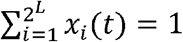, as it is expected. In the fixed environment ***e***(*t*) and in the absence of HGT, *p*_*HGT*_ = 0, we recover the discrete-time version of the well-known replicator-mutator equation (46–48).

Using (5), we can find each term of the transition matrix *A*_*ij*_

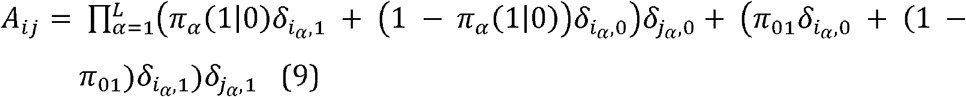

Here, *δ*_*i,k*_ if *i = k* and *δ*_*i,k*_ = 0 if *i ≠ k* is the Kronecker delta. *π* is given by (6).

In (8), the mean fitness of the population, 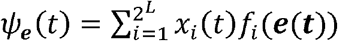 describes the growth of the total population in the environment ***e***(*t*). However, it varies stochastically due to the environmental variations. To quantify the long-term growth of the population we discuss the following quantity

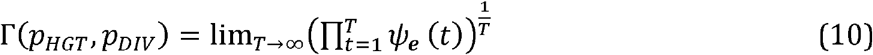

Note that if environment is fixed, that is ***e***(*t*) = *const*, then the long-term growth of the population is defined by the mean fitness of the population *ψ*_*e*_(*t*).

In (10), we explicitly mentioned *p*_*h*_ and *p*_*ad*_ in the argument of the long-term fitness function because the optimal values of these quantities are found from maximization of *Γ*(*p*_*HGT*_, *p*_*DIV*_) over *p*_*HGT*_, *p*_*DIV*_ ∈[0,1], for the given values of viruses statistics, death probabilities, reproduction costs and gene lose. The maximization algorithm is provided in SI section V.

## Data Availability

No data was used in this work.

## Acknowledgements

The authors thank Koonin group members for many helpful discussions. The authors’ research is supported by the Intramural Research Program of the National Institutes of Health (National Library of Medicine). S.G.B. was also supported by the Higher Education and Science Committee of RA (Research Program 24IRF/2-1C001). This work utilized the computational resources of the NIH HPC Biowulf cluster (http://hpc.nih.gov).

## Author contributions

S.G.B., Y.I.W. and E.V.K. conceptualized the model; S.G.B. developed the model; S.G.B., S.K.G., Y.I.W and E.V.K. analyzed the results; S.G.B. and E.V.K wrote the manuscript that was edited and approved by all authors.

## Notes

### Competing Interest Statement

The authors have declared no competing interest.

