## Supporting Information for "Evolution of antivirus defense in prokaryotes depending on the environmental virus prevalence and virome dynamics"

### Supplementary Information

#### I. SHORT REVIEW OF TWO-STATE MARKOV CHAIN

In the main text, we indicate that viruses are described by their probability of presence and characteristic dynamics times, $\mu$ and $\tau$, respectively. Here, we outline the connections between these quantities and the appearance and disappearance probabilities, $q$ and $p$, respectively, for one virus case. Consider the following Markov chain

$$P = \begin{pmatrix} 1-p & q \\ p & 1-q \end{pmatrix} \tag{1}$$

where $P$ is the transition matrix. The first row shows the transition probabilities to state 1 from 1 and 0. That is, with probabilities $1-p$ and $q$ a currently present and absent virus will stay or appear in the environment, respectively. The matrix $P$ is stochastic, that is, $\sum_j P_{ij} = 1$ and $P_{ij} > 0$.

The eigenvalues are $\lambda_1 = 1$ and $\lambda_2 = |1-p-q|$. The row vector $(1,1)$ is a left-eigenvector of the matrix $P$, that corresponds to the eigenvalue $\lambda_1 = 1$. The transition matrix has two right and left eigenvectors that are orthonormal. That is, there are two pairs of right (column) and left (row) that satisfy the following relations.

$$\mathbf{l}_i P = \lambda_i \mathbf{l}_i \tag{2}$$

$$P\mathbf{r}_i = \lambda_i \mathbf{r}_i \tag{3}$$

where $i = 1,2$. These eigenvectors are $\mathbf{r}_1 = (\mu, 1-\mu)^T$, $\mathbf{r}_2 = (1,-1)^T$, $\mathbf{l}_1 = (1,1)$ and $\mathbf{l}_2 = (1-\mu, -\mu)$, where $\mu = \frac{q}{p+q}$, that is, the virus presence probability. Note that $\mathbf{l}_i\mathbf{r}_j = \delta_{ij}$, where $\delta_{ij}$ is the Kroenecker delta.

The matrix $P$ is diagonalizable, that is, $P = U^{-1}\Lambda U$, where $\Lambda$ is a diagonal matrix, with $\lambda_1$ and $\lambda_2$ (eigenvalues of $P$) on the diagonal. The matrix $U$ has the left eigenvectors $\mathbf{l}_i$ as rows, while $U^{-1}$ has right eigenvectors, $\mathbf{r}_i$, as columns with the obvious relation $U^{-1}U = I$. Then, the $t$-step transition matrix is equal to $P^t = U^{-1}PU\ U^{-1}PU... = U^{-1}P^tU$. Now let us suppose that initially the virus is present, that is, the initial state of the environment is $e_0 = (1,0)^T$. Let us then find the state of the environment at time $t$.

$$e_t = P^t e_0 = \begin{pmatrix} \mu + (1-\mu)\lambda_2^t \\ 1-\mu-(1-\mu)\lambda_2^t \end{pmatrix} \tag{4}$$

Recalling that $\lambda_2 = |1-p-q| \leq 1$, we get that the smaller $\lambda_2$ is the faster the environment converges to its stationary state $\mathbf{r}_1 = (\mu, 1-\mu)^T$, and accordingly, the faster any correlations between the environmental states disappear. We introduce $\tau$ as a characteristic time that is connected to $\lambda_2$ in the following way $e^{-t/\tau} = {\lambda_2}^t$. Thus, greater $\lambda_2$ implies greater $\tau$, and environment converges to the stationary state slower.

#### II. EXPECTED VALUE OF THE FITNESS

Let us find the expected value of the fitness of a cell with an immune repertoire $\mathbf{s}$ over the environmental fluctuations. The fitness of a cell with immune repertoire $\mathbf{s}$ in the environment $\mathbf{e}$ is given by the following expression

$$f_{\mathbf{s}} = (1-c_{DIV})^{p_{DIV}}(1-d_0)\prod_{\alpha=1}^{L}(1-c_\alpha s_\alpha)(1-d_\alpha e_\alpha(1-s_\alpha)) \tag{5}$$

Note, that $e_\alpha = 0, 1$, $\alpha = 1, ..., L$. The expected value of $f_{\mathbf{s}}$ is calculated as follows

$$\overline{f_s} = \sum_{e_1=0,1}\sum_{e_2=0,1}..\sum_{e_L=0,1}(1-c_{DIV})^{p_{DIV}}(1-d_0)\prod_{\alpha=1}^{L}(1-c_\alpha s_\alpha)(1-d_\alpha e_\alpha(1-s_\alpha))\Pr(\mathbf{e}) =$$

$$= (1-c_{DIV})^{p_{DIV}}(1-d_0)\sum_{e_1=0,1}\sum_{e_2=0,1}..\prod_{\alpha=1}^{L-1}(1-c_\alpha s_\alpha)(1-d_\alpha e_\alpha(1-s_\alpha))\Pr(e_\alpha)\sum_{e_l=0,1}(1-c_L s_L)(1-d_L e_L(1-s_L))\Pr(e_L)$$

$$= (1-c_{DIV})^{p_{DIV}}(1-d_0)\prod_{\alpha=1}^{L}(1-c_\alpha s_\alpha)(1-\mu_\alpha d_\alpha(1-s_\alpha)) \quad (6)$$

In the first line $\Pr(\mathbf{e})$ is the probability of any configuration of environment, that is, any of $2^L$ states. In the second line, we apply the independence assumption of virus dynamics $\Pr(\mathbf{e}) = \prod_{\alpha=1}^{L}\Pr(e_\alpha)$. Here, we also use the fact that fitness function is a product of independent terms as well. Note that $\Pr(e_\alpha) = 1$ is equal to $\mu_\alpha$. Then, $\sum_{e_l=0,1}(1-c_\alpha s_\alpha)(1-d_\alpha e_\alpha(1-s_\alpha))\Pr(e_\alpha) = (1-c_\alpha s_\alpha)(\mu(1-d_\alpha(1-s_\alpha)+(1-\mu)(1-0)) = (1-c_\alpha s_\alpha)(1-\mu_\alpha d_\alpha(1-s_\alpha)$.

Now, let us compare the expected fitness values of two cells that have the same immune repertoire except at position $\gamma$, that is $s_\gamma = 1$ and $s'_\gamma = 0$. Then we get

$$\frac{\overline{f_{\mathbf{s}}}}{\overline{f_{\mathbf{s}'}}} = \frac{1-c_\gamma s_\gamma}{1-\mu_\gamma d_\gamma(1-s'_\gamma)} = \frac{1-c_\gamma}{1-\mu_\gamma d_\gamma} \quad (7)$$

From (7), it follows that $\frac{\overline{f_{\mathbf{s}}}}{\overline{f_{\mathbf{s}'}}} > 1$ if $\mu_\gamma > \frac{c_\gamma}{d_\gamma}$. From the last condition, it follows that in the symmetric case, that is, $\mu_\alpha = \mu$, $c_\alpha = c$ and $d_\alpha = d$ for all $\alpha = 1, ..., L$, either naive or fully immune cells have the maximum average fitness, that is, the cells with $s_\alpha = 0$ and $s_\alpha = 1$ , $\alpha = 1, ..., L$, respectively. In the asymmetric case, cells with neither full nor empty immune repertoires may be the most fittest on average, depending on the environment.

#### III. MULTIPLICATIVE V.S. ADDITIVE COST OF IMMUNITY

In the main text, it is stated that multiplicative fitness reduction provides greater fitness value than additive reduction for the given set of the costs of the defense genes $\{c_\alpha > 0\}$, such that $\sum_{\alpha=1} c_\alpha < 1$. Note that, if the last condition does not hold, then, obviously multiplicative reduction will be smaller because the $\prod_{\alpha=1}^{L}(1-c_\alpha) > 0$.

To prove the statement, let us first observe that it holds for the case of two costs. Indeed, in the case, we get $(1-c_1)(1-c_2) = 1-c_1-c_2+c_1c_2 > 1-c_1+c_2$. Now let us assume that the statement holds for some $k$, and let's prove that it is also true for $k+1$, that is

$$\prod_{i=1}^{k+1}(1-c_\alpha) > 1-\sum_{i=1}^{k+1}c_\alpha \quad (8)$$

$$(\prod_{i=1}^{k}(1-c_\alpha) - (1-\sum_{i=1}^{k}c_\alpha)) + c_{k+1}(1-\prod_{i=1}^{k}(1-c_\alpha)) > 0 \quad (9)$$

The last equality is valid by the assumption that the statement is true for $k$, and all costs are positive $c_\alpha > 0$.

#### IV. LONG TERM GROWTH AND COST OF ADAPTIVE IMMUNITY

In the main text, it is pointed out that increasing the cost of DIV immunity, $c_{DIV}$, decreases the maximum possible value of the optimal probability of defense acquisition via this rout $p_{DIV}$. Here, we will derive the maximum possible long-term fitness and compare it with suboptimal growth of the population, to characterize the interplay between the cost and efficiency of immunity acquired via DIV.

Let us start from the definition of the long-term growth fitness of the population, under the assumption that in each environment the population reaches the maximum possible fitness

$$\Gamma = \big(\prod_{t=1}^{T} \psi_{\mathbf{e}}(t)\big)^{1/T} \leq \big(\prod_{t=1}^{T} \max[f_1(\mathbf{e}(t)), ..f_{2^L}(\mathbf{e}(t))]\big)^{1/T} \tag{10}$$

The last assumption, obviously does not hold given the gene loss and defense acquisition. Indeed, suppose that $f_{\mathbf{s}}$ is the maximum fitness in the environment $\mathbf{e}(t)$; then, due to the transition probabilities from $\mathbf{s} \to \mathbf{s}'$, where $\mathbf{s} \neq \mathbf{s}'$, the population will not be homogeneous in that environment. Therefore, the approach is still useful to elucidate the behavior of $p_{DIV}$ depending on the cost of DIV, $c_{DIV}$.

From (10), it follows that

$$\begin{aligned}\ln\Gamma = \frac{1}{T}\sum_{t=1}^{T} \ln\psi_{\mathbf{e}}(t) \leq \ \frac{1}{T}\sum_{t=1}^{T} \ln(\max[f_1(\mathbf{e}(t)), ..f_{2^L}(\mathbf{e}(t))]) = \\ = E\big[\ln(\max[f_1(\mathbf{e}(t)), ..f_{2^L}(\mathbf{e}(t))]\big]\end{aligned} \tag{11}$$

Assuming that $T \to \infty$ and noticing that neither the population nor the virus dynamics have absorbing states, we can approximate the sum in the equation by its average over all possible environmental fluctuations. Let us first consider one virus case, with the assumption that $c < d$. In this case, the environment is $e = 0, 1$. Denoting the fitness of naive and immune cells by $f_1$ and $f_2$, respectively, we get for the maximum fitness in each environment

$$\max[f_1, f_2] = \begin{cases} (1 - c_{DIV})^{p_{DIV}}(1 - d_0), & \text{if } e = 0 \text{ w.p. } 1 - \mu \\ (1 - c_{DIV})^{p_{DIV}}(1 - d_0)(1 - c), & \text{if } e = 1 \text{ w.p. } \mu \end{cases} \tag{12}$$

Then, the average of the logarithm of the maximum fitness is $E[\ln[\max[f_1, f_2]]] = p_{DIV}\ \ln(1 - c_{DIV}) + \ln(1 - d_0) + \mu\ln(1 - c)$

Thus, we get the following expression for the maximum long term growth of the population

$$\ln\Gamma_{\max} = p_{DIV}\ \ln(1 - c_{DIV}) + \ln(1 - d_0) + \mu\ \ln(1 - c) \tag{13}$$

The suboptimal growth in this case corresponds to the long-term growth of the naive population, that is, assuming $p_{DIV} = 0$. The suboptimal growth is given by

$$\ln\Gamma_{\text{sub}} = \ln(1 - d_0) + \mu\ \ln(1 - d) \tag{14}$$

Comparing (13) and (14) we find the threshold value of $p_{DIV}$ for which the long-term maximum becomes equal to the suboptimal value

$$\tilde{p}_{DIV} = \mu\frac{\ln(1 - d) - \ln(1 - c)}{\ln(1 - c_{DIV})} \tag{15}$$

Thus, $\ln\Gamma_{\max} > \ln\Gamma_{\text{sub}}$ for $p_{DIV} \in (0, \tilde{p})$, if $\tilde{p} > 1$, then $\ln\Gamma_{\max} > \ln\Gamma_{\text{sub}}$ in the whole region $p_{DIV} \in (0, 1)$. From (15) it follows that $\frac{\partial\tilde{p}_{DIV}}{\partial d} > 0$ and $\frac{\partial\tilde{p}_{DIV}}{\partial c}$, $\frac{\partial\tilde{p}_{DIV}}{\partial c_{DIV}} < 0$. That is, increasing the probability of death caused by the viruses increases the threshold value of the $p_{DIV}$ below which the maximum fitness is still greater than the suboptimal one. Conversely, increasing the cost of defense and extra cost of DIV mechanism decreases the threshold value. Furthermore, it follows that increasing virus prevalence in the environment increases $\tilde{p}_{DIV}$, and $\tilde{p}_{DIV} = \mu$ when $c_{DIV} = \frac{d-c}{1-c}$.

In the case of three pathogens, with the same cost of defenses and death probabilities, we get for the same expression (15) for the threshold value, still assuming that the suboptimal fitness corresponds to a naive population, except that $\mu \to \mu_1 + \mu_2 + \mu_3$. Thus, $\tilde{p}_{DIV}$ increases when there are more viruses in the environment, that is, more room for optimization.

#### V. OPTIMIZATION ALGORITHM FOR FINDING THE OPTIMAL VALUES OF HGT AND ADAPTIVE IMMUNITY

We need to maximize the long-term growth of the population over the parameters $(p_{HGT}, p_{DIV})$ to find the optimal values of the probabilities of defense acquisition through HGT and DIV. Note that for any pair of values $(p_{HGT}, p_{DIV})$, $\Gamma(p_{HGT}, p_{DIV})$ in each run is random because of stochastic changes in the environment. So, we need to find the optimal pairs of parameters $(p_{HGT}, p_{DIV})$, which maximize stochastic function. The provided algorithm is adopted from [1], with several modifications. The optimization is over simplex $[0, 1] \times [0, 1]$, with the following steps

1. Initialize the step length and number of samples $\epsilon, N$.
2. Initialize $(p_{HGT}, p_{DIV})$ in the simplex $[0, 1] \times [0, 1]$. Construct the Moore neighborhood for the point $(p_{n,HGT}, p_{n,DIV}) = \{(p_{HGT} + a, p_{DIV} + b);\ a, b = \{-\epsilon, 0, \epsilon\}\}$ but $a = b = 0$.
3. For each pair in the neighborhood of $(p_{j,HGT}, p_{j,DIV})$, $j \in$ neighborhood, sample $\Gamma_N(p_{j,HGT}, p_{j,DIV})$, $N$ times. In this step, for each neighbor we get a list $\langle\Gamma_N(p_{j,HGT}, p_{j,DIV})\rangle$ where each element represents a value of the long-term growth for the given pairs of the parameters.
4. Select the pairs $(p_{j,HGT}, p_{j,DIV})$ from the neighbors that have different than $(p_{HGT}, p_{DIV})$ population mean, inferred from $\langle\Gamma(p_{j,HGT}, p_{j,DIV})\rangle$ by unequal variance t-test with p-value equal to 0.05.
5. From all neighbors that have different means select the one that has the maximum population mean. That is select a pair $(p_{k,HGT}, p_{k,DIV}) = \text{argmax}\,\{E[\Gamma_N(p_{k,HGT}, p_{k,DIV})]; k \in S\}$, where $S = \{j \in \text{neighborhood}; E[\langle[\Gamma_N(p_{j,HGT}, p_{j,DIV})\rangle] \neq E[\langle\Gamma_N(p_{HGT}, p_{DIV})\rangle]\} \cap (p_{HGT}, p_{DIV})$.
6. Update the current point $(p_{HGT}, p_{DIV}) \leftarrow (p_{k,HGT}, p_{k,DIV})$ and go to step 1.
7. If neither of the neighbors pass 4, then go to 1 and update $\{\epsilon, N\} = \{\epsilon/2, 2N\}$.
8. If $\epsilon < \epsilon_0$, then return the current value $(p_{HGT}, p_{DIV})$.

The parameters in the algorithm are set to $\epsilon = 0.01$, $\epsilon_0 = 0.005$ and $N = 100$. Note, that the algorithm provided above finds local maximum, instead of global maximum. However; when the fitness function has one peak, then the algorithm finds it. This is the case for extreme values of pathogen presence and characteristic times. In the long-term fitness has one but flat peak, then it finds a value on the flat peak.

#### VI. NOTES ON POPULATION DYNAMICS: AVERAGED OVER ENVIRONMENT.

In the main text we claim that in the HGT-dominated region of defense acquisition process, that is $p^*_{HGT} = 1$ and $p^*_{DIV} \approx 0$, we observe that the fractions of cells carrying different immune repertoires show universal behavior defined only by the gene loss probability, $\pi_{01}$. Here we consider the genesis of the observed effect.

First, let us observe that for small characteristic times $\tau \leq 1$, the population dynamics can be approximated by

$$x_i(t+1) = E\left[\frac{\sum_{j=1}^{2^L} x_j(t) A_{ij}\big(\mathbf{x}(t), \mathbf{e}(t)\big) f_j(\mathbf{e}(t))}{\sum_{j=1}^{2L} x_j(t) f_j(\mathbf{e}(t))}\right] \tag{16}$$

where the average is over the environmental fluctuations. In the HGT-dominated region, the matrix $A$, describing the transitions of cells between different immune repertoires, becomes weakly dependent on the environment $\mathbf{e}(t)$. Indeed, the dependency of the matrix $A$ on the state of the environment is due to the defense acquisition via DIV $\pi_\alpha(1|0, \mathbf{x}(t), \mathbf{e}(t) = 1 - (1 - p_{DIV} e_\alpha(t))\big(1 - p_{HGT}(\sum_{k=1}^{2^L} x_k(t)\mathbf{s}_k)_\alpha\big)$, which becomes a weakly dependent on the environment in the HGT-dominated region due to the $p^*_{DIV} \approx 0$. We will assume that $p^*_{DIV} = 0$, although, strictly speaking, in this case, cells with different immune repertoires will not appear from naive population.

Therefore, the fitnesses of cells in (16) are still environment-dependent quantities. Let us consider the dynamical system (16) around the averages of the fitnesses, that is around $\overline{f} = \{\overline{f_1}, \overline{f_2}, ..., \overline{f_{2^L}}\}$, enumeration is according to the fractions $x_i, i = 1, , 2^L$.

Taylor expanding the right hand-side of (16) around the mean fitnesses, we get the following dynamical system

$$
\begin{aligned}
x_i(t+1) &= \sum_{j=1}^{2^L} x_j A_{ij}\Big(\frac{\overline{f_j}}{\sum_{j=1}^{2L} x_j \overline{f_j}} + E\big[\sum_l \frac{\partial}{\partial f_l}\frac{f_j}{\sum_k x_k f_k}|_{\overline{f}}(f-\overline{f_l})\big] + \\
&+\frac{1}{2}E\big[\sum_{l,m}\frac{\partial}{\partial f_m}\frac{\partial}{\partial f_l}\frac{f_j}{\sum_k x_k f_k}|_{\overline{f}}(f_m-\overline{f_m})(f_l-\overline{f_l})\big] + ..\Big) = \\
&= \frac{\sum_{j=1}^{2^L} x_j A_{ij}}{\sum_k x_k \overline{f_k}}\Big(\overline{f_j} + E\big[\sum_{l,m}(f_l-\overline{f_l})\big(\frac{\overline{f_j} x_m x_l}{(\sum_k x_k \overline{f_k})^2} - \frac{\delta_{jm}x_l+\delta_{jl}x_m}{2\sum_k x_k \overline{f_k}}\big)(f_m-\overline{f_m})\big]\Big) \\
&= \frac{\sum_{j=1}^{2^L} x_j A_{ij}\overline{f_j}}{\sum_k x_k \overline{f_k}}
\end{aligned}
\tag{17}
$$

The term containing averaging over environmental fluctuations is $E[] \sim O(\text{var} f_j)$ in the considered case, $x_l, x_m \sim x$ and $f_j, f_k \sim f$, then the expectation is $\sim O(\text{var} f)$. Then the contribution from the higher order terms is small compared to $\overline{f}$

$$
O(\frac{\text{var}\overline{f}}{\overline{f}}) \sim \prod_{\alpha=1}^{L} 1-\mu_\alpha d_\alpha(1-s_\alpha) < 1 \tag{18}
$$

if $\mu_\alpha d_a > 0$. We numerically checked the solutions obtained from (16) and (17), for $p^*_{HGT} = 1$ and varying $p_{DIV}$ in (16), but keeping $p_{DIV} \approx 0$ in (17). Fig.1a shows the euclidean distance between the composition of the population in the equilibria obtained from (16), $\mathbf{x}_{sol}$ and (17), $\mathbf{x}_{apr}$.For $p_{DIV} \approx 0$, the two solutions agree well, but increasing $p_{DIV}$ leads to an increasing discrepancy between the solutions. Indeed, for $p_{DIV} > 0$ the correlations between the matrix $A$ and fitnesses cannot be neglected, hence the increasing discrepancy. We further compared the optimal values of HGT and DIV found from (17) that maximizes the population fitness $\sum_k x_k f_k$ in equilibrium, $(p_{HGT}, p_{DIV})_{apr}$, with the values obtained from the stochastic optimization algorithm $(p_{HGT}, p_{DIV})_{st.op}$, see Fig.1b. As in the main text we choose $\{\mu_1, \mu_2, \mu_3\} = \{3/2\mu, \mu/2, \mu\}$. As seen in the figure, both methods produce the same optimal values for high viral load $\mu$, that is in the HGT-dominated region. However, the approach developed in this section fails for small viral load $\mu = 0.3$ and longer characteristic times, because in both cases the optimal values of defense acquisition probabilities are out of HGT-dominated region. In the former case, it is due to the small virus prevalence (but close to $\mu_{thr}$), where $(p_{HGT}, p_{DIV})_{apr} = (0, 0)$ but stochastic optimization still yields non-zero values of $(p_{HGT}, p_{DIV})_{st.op}$. In the latter case, $p^*_{DIV} \neq 0$ in the environments with longer characteristic times. Indeed, for longer characteristic times of pathogen dynamics (16) is not applicable, and hence the approach developed thereafter.

#### VII. ANALYSIS OF THE HGT-DOMINATED REGION.

In the HGT-dominated region, the frequencies of cells with different immune repertoires in equilibrium are given by

$$
x^*_i = \pi_{01}^{L-|\mathbf{s}_i|}(1-\pi_{01})^{|\mathbf{s}_i|} \tag{19}
$$

where, for future convenience, we denote $|\mathbf{s}_i| = \sum_{\alpha=1}^{L} s_{i,\alpha}$ that is, the number of defense genes in the repertoire $\mathbf{s}_i$. Now, let us calculate the probabilities of the acquisition of defense genes for any virus $\alpha = 1, ..., L$ in equilibrium (19). Recalling the definition of $\pi_\alpha(1|0)$ and taking into account that the optimal values of the defense acquisition process through HGT and DIV are $p^*_{HGT} = 1$ and $p^*_{DIV} = 0$ in the HGT-dominated region, respectively, we get

$$
\pi_\alpha(1|0) = \big(\sum_{k=1}^{2^L} x^*_k \mathbf{s}_k\big)_\alpha \tag{20}
$$

Note that in the sum $x^*_k$ is a scalar and $\mathbf{s}_k$ is a vector, thus the right-hand side is a convex combination of immune repertoires $\mathbf{s}_k,\ k = 1, ..., 2^L$, and the $\alpha$-th element of the resulting vector shows the probability of acquisition of

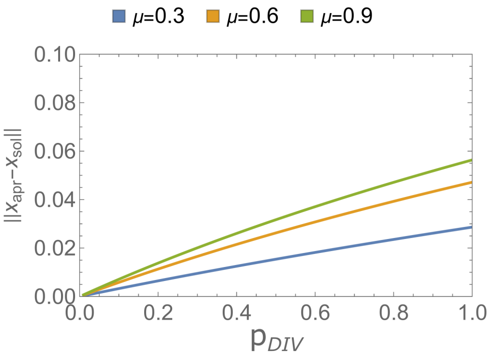


(a)

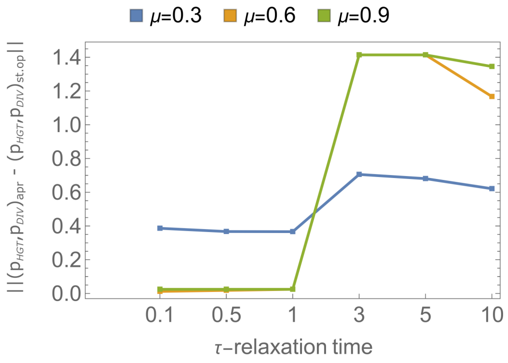


(b)

Supplementary Figure 1: Comparison between different approaches. a) Euclidean distance between the composition of the population obtained from (16) and (17), where we neglected higher-order terms. Here, $p_{HGT} = 1$. b) Euclidean distance between the optimal values of HGT and adaptive immunity obtained from optimization of the mean fitness of the population at equilibrium found from (17) and stochastic optimization algorithm discussed in the previous section and used in the manuscript. The model parameters are $\{\mu_1, \mu_2, \mu_3\} = \{3/2\mu, \mu/2, \mu\}$, $\tau_1 = \tau_2 = \tau_3 = 0.1$, $c_\alpha = 0.3$, $d_\alpha = 0.7$, $c_{DIV} = 0.1$, $d_0 = 0.1$.

defense against the virus $\alpha$. Note that for each virus $\alpha = 1, ..., L$ there are $2^{L-1}$ immune repertoires that could serve as a donor for HGT, that is, in all these immune repertoires $s_\alpha = 1$. Then, among all of these $2^{L-1}$ repertoires, there is only one immune repertoire with $|\mathbf{s}| = 1$, the one in which $s_\alpha = 1$, whereas all other elements are zeros $s_{\gamma\neq\alpha} = 0,\ \gamma = 1, .., L-1$. Similarly, there are $C_{L-1}^{L-2}$ immune repertoires with $|\mathbf{s}| = 2$, one defense gene is in $\alpha$ and one in the remaining $L-1$ positions, which corresponds to the number of different combinations, immune repertoires, that have $L-2$ zeros in $L-1$ places. Thus, for (20) we obtain

$$\pi_\alpha(1|0) = C_{L-1}^{L-1}\pi_{01}^{L-1}(1-\pi_{01}) + C_{L-1}^{L-2}\pi_{01}^{L-2}(1-\pi_{01})^2 + ... + C_{L-1}^{0}(1-\pi_{01})^L =$$
$$= (1-\pi_{01})\sum_{k=0}^{L-1} C_{L-1}^{k}(\pi_{01} + (1-\pi_{01}))^k = 1 - \pi_{01}. \tag{21}$$

Thus, in the equilibrium (19), the probability of defense acquisition against any pathogen is the same and equal to $1-\pi_{01}$. For the transition matrix $A$, we obtain

$$A_{ij}(\mathbf{x}^*) = \prod_{\alpha=1}^{L}(\pi_\alpha(1|0)\delta_{i_\alpha,1} + (1-\pi_\alpha(1|0))\delta_{i_\alpha,0})\delta_{j_\alpha,0} + (\pi_{01}\delta_{i_\alpha,0} + (1-\pi_{01})\delta_{i_\alpha,1})\delta_{j_\alpha,1} =$$
$$= \prod_{\alpha=1}^{L}((1-\pi_{01})\delta_{i_\alpha,1} + \pi_{01}\delta_{i_\alpha,0})(\delta_{j_\alpha,0} + \delta_{j_\alpha,1}) = \pi_{01}^{L-|\mathbf{s}_i|}(1-\pi_{01})^{|\mathbf{s}_i|} = x_i^* \tag{22}$$

here we used $\delta_{j_\alpha,0} + \delta_{j_\alpha,1} = 1$, for any $\alpha$. Thus, the transition probabilities become dependent only on resulting immune repertoire $A_{ij} = A_i$, furthermore, all elements in each row $i$ is the same.

We checked numerically that the equilibrium (19) is stable, whenever it appears. Indeed, let us consider the equivalent continuous-time version of the dynamical system (17),

$$\frac{dx_i}{dt} = \sum_j x_j A_{ij}\overline{f_j} - \sum_j x_j\overline{f_j}x_i \tag{23}$$

Indeed, the state (19) is an equilibrium of (23). In the equilibrium state, the right-hand side vanishes $\sum_j x_j^* A_{ij}(\mathbf{x}^*)\overline{f_j} - \sum_j x_j^*\overline{f_j}x_i^* = 0$, recalling that $A_{ij}|_{\mathbf{x}^*} = A_i|_{\mathbf{x}^*}$, we get $x_i^* = A_i|_{\mathbf{x}^*}$. The matrix elements of the Jacobian are

$$\frac{\partial}{\partial x_n}\frac{dx_i}{dt} = \overline{f_n}(A_{in} - x_i)|_{\mathbf{x}^*} - \delta_{in}\sum_k x_k^*\overline{f_k} + \sum_j x_j \partial_n A_{ij}(\mathbf{x}^*)\overline{f_j} \tag{24}$$

Recalling (22), the Jacobian matrix becomes

$$\mathbf{J}|_{\mathbf{x}^*} = -Diag[\sum_k x_k\overline{f_k}]|_{\mathbf{x}^*} + \mathbf{x}\frac{\partial A(\mathbf{x})}{\partial \mathbf{x}}\mathrm{Diag}[\overline{f_k}]|_{\mathbf{x}^*} \tag{25}$$

The last term is difficult to evaluate, but checking the eigenvalues of the Jacobian in the equilibrium (19) gives that the largest eigenvalue by modulo is equal to to the mean fitness of the population $\sum_k x_k^* f_k$.

In equilibrium (19), the long-term fitness of the population, which is the same as the mean fitness of the population, is equal to

$$\Gamma^* = \sum_k x_k^*\overline{f_k} = (1-d_0)\sum_{j=1}^{2^L} \pi_{01}^{L-|\mathbf{s}|_j}(1-\pi_{01})^{|\mathbf{s}|_j}\prod_{\alpha=1}^{L}(1-c_\alpha s_{j,\alpha})(1-\mu_\alpha d_\alpha(1-s_{j,\alpha})) \tag{26}$$

For identical viruses, that is $\mu_\alpha = \mu$, $c_\alpha = c$ and $d_\alpha = d$, (26) simplifies to

$$\Gamma^*_{sym} = (1-d_0)\big(1-c+\pi_{01}(c-\mu d)\big)^L \tag{27}$$

Note that equilibrium (19) occurs when $c < \mu d$.

#### VIII. SUPPLEMENTARY FIGURES

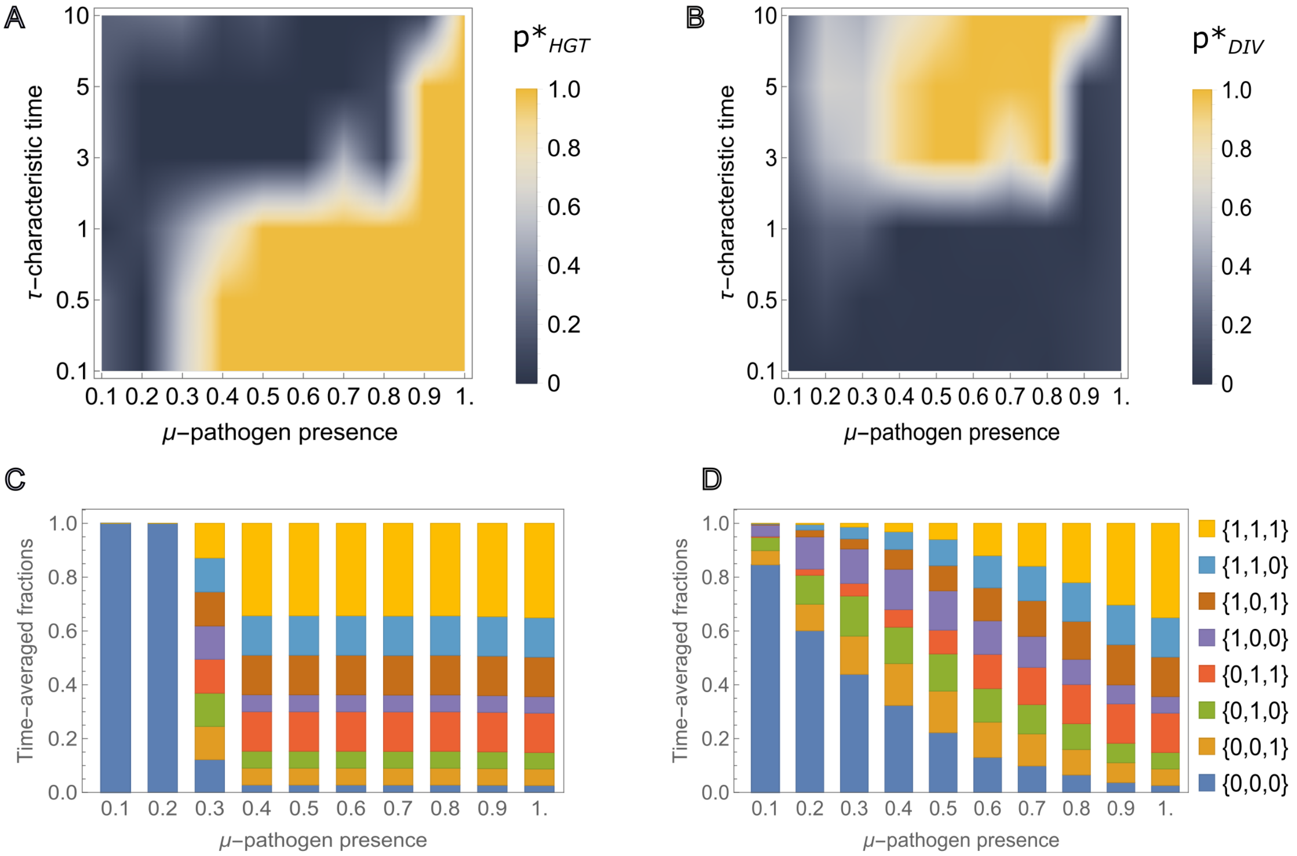


Supplementary Figure 2: Phase space and population composition for equal probabilities of virus presence and characteristic times.
A) and B), optimal values of the probability of defense acquisition through HGT and DIV, respectively, for various viral load in the environment.
C) and D), Composition of the population averaged over time for $\tau = 0.1$ and $\tau = 10$, respectively.
Here we assume that all viruses have the same characteristic times and virus presence probability $\tau$ and $\mu$, respectively. The remaining model parameters are $c_\alpha = 0.3$, $d_\alpha = 0.7$, $c_{DIV} = 0.1$, $d_0 = 0.1$.

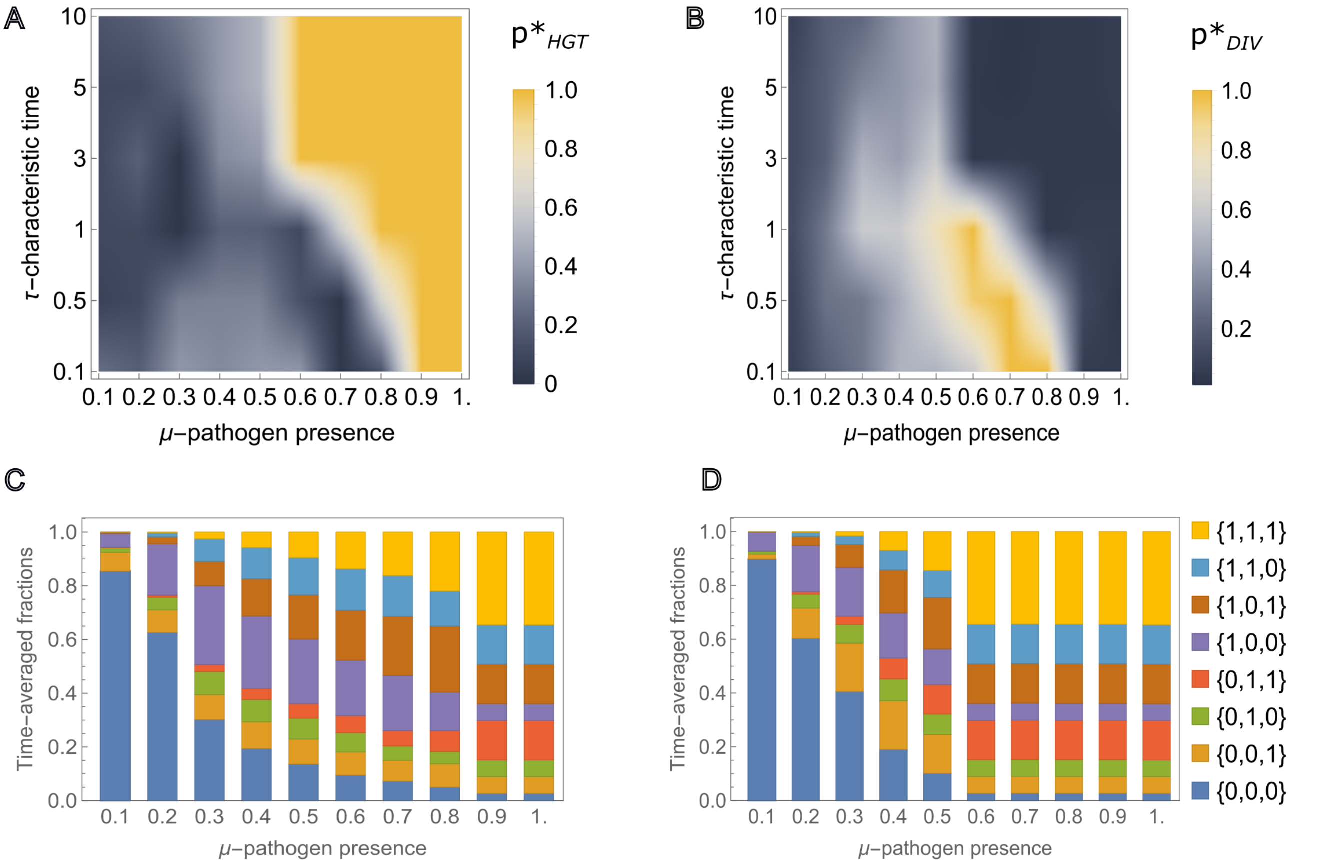


Supplementary Figure 3: Phase space and population composition for unequal probabilities of virus presence and characteristic times.
A) and B), optimal values of the probability of defense acquisition through HGT and DIV, respectively, for various virus presence in the environment.
Here we assume that less frequent virus has fixed characteristic time $\tau_2 = 1$. The characteristic times of remaining two viruses taken from the list $\tilde{\tau} = \{0.1, 0.5, 1, 3, 5, 10\}$, such that $\tau \equiv \tau_1 = \tilde{\tau}_j$ and $\tau_3 = \tilde{\tau}_{-j}$, where $j = 1, ...6$ shows the index of the elements in $\tilde{\tau}$, $-j$ means counting from the last element of $\tilde{\tau}$. That is, the remaining two viruses are always in pairs, where one of them has smaller characteristic times than the other one.
C) and D), Composition of the population averaged over time for $\{\tau_1, \tau_2, \tau_3\} = \{0.1, 1, 10\}$ and $\{\tau_1, \tau_2, \tau_3\} = \{10, 1, 0.1\}$, respectively.
The remaining model parameters are $\{\mu_1, \mu_2, \mu_3\} = \{3\mu/2, \mu/2, \mu\}$, $c_\alpha = 0.3$, $d_\alpha = 0.7$, $c_{DIV} = 0.1$, $d_0 = 0.1$.

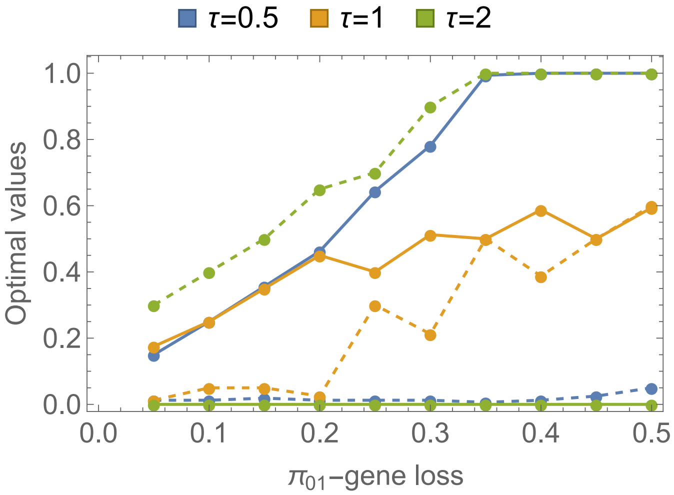


Supplementary Figure 4: Dependence of optimal values of HGT (full) and DIV (dashed) for various gene loss probabilities and characteristic times of pathogens.
The probabilities of pathogens presence are $\{\mu_1, \mu_2, \mu_3\} = \{3\mu/2, \mu/2, \mu\}$ with $\mu = 0.4$. The characteristic times are the same for all pathogens, $\tau_\alpha = \tau$. The remaining model parameters are the same as in Fig.2.

[1] Mayer, Andreas, et al. "Diversity of immune strategies explained by adaptation to pathogen statistics." Proceedings of the National Academy of Sciences 113.31 (2016): 8630-8635.
